# Missense3D-PTMdb an interactive web resource to explore and visualize genetic variants and post-translational modifications sites (PTMs) using AlphaFold 3D models

**DOI:** 10.1101/2025.07.22.666170

**Authors:** Haotian Zhao, Ryan Pye, Grace Walker, Wendy Tran, Olivia Simmonds, Ifigenia Tsitsa, Suhail Islam, Gordon Hanna, Alessia David

**Affiliations:** Centre for Bioinformatics, Department of Life Sciences, Imperial College London, London SW7 2AZ UK

## Abstract

Only a fraction of the >11 million missense variants identified in the human population has a known damaging or tolerated clinical impact. Post-translational modifications (PTMs), such as phosphorylation, glycosylation and ubiquitination, are key regulators of protein function and structure, and are critical for protein localisation, stability and interactions with other molecules. The ability of a protein to undergo PTMs, is subject to a correctly folded protein structure, and the recognition and binding of enzymes to specific amino acid motifs in close proximity to residues that undergo PTMs [PTM residue]. AlphaFold models provide an unprecedented opportunity to perform sequence-structure mapping of variants, which are in close linear or spatial proximity to a PTM site and should have their impact on protein function experimentally investigated.

We present Missense3D-PTMdb, a “one-stop-shop” interactive web tool that provides a user-friendly sequence-structure mapping of 20,235 human proteins to 11,544,303 naturally occurring human missense variants, 203,775 PTM sites and their neighbours in sequence and 3D structure space using AlphaFold generated 3D models of the human proteome. Additionally, the sequence-structure mapping tool allows visualization and exploration of any human variant not currently stored in the database. Missense3D-PTMDb is freely available at https://missense3d.bc.ic.ac.uk/ptmdb.

## Introduction

Large-scale genome projects are generating an extensive amount of genetic variation data yet the characterisation and prioritisation of variants that may affect phenotype and cause disease remains a significant challenge. Post-translational modifications (PTMs), such as phosphorylation, glycosylation, methylation, and ubiquitination are key regulators of protein function and structure. PTMs are critical for protein localisation, stability and interactions with other molecules ^1, 2^. Moreover, PTMs expand the functional diversity of proteins beyond what is already encoded by the coding sequence, enabling the rapid generation of phenotypic complexity and divergence ^3^. Disruption of a protein’s ability to undergo PTMs can impair numerous cellular processes, including transcriptional regulation, cell metabolism and differentiation ^4^. One fundamental step towards understanding the role of genetic variants in disrupting protein function, is the ability to map and inspect their position, relative to PTM sites. This includes identifying variants that occur directly on residues that undergo post-translational modifications (PTM-residues), as well as those located in close linear and/or spatial proximity to PTM-residues, which may perturb PTMs by disrupting recognition motifs and/or by influencing the structural stability of the protein ^4, 5^.

At present, no single comprehensive resource provides 3D structural mapping of PTMs, human variants and disease information ^6^. The recent development of AlphaFold, which produces highly accurate 3D protein structure models of the human proteome, ^7, 8^ provides an unmissable opportunity to fill this gap. Building upon our existing Missense3D Portal for the structural characterization of missense variants using 3D structures ^9^, we have developed Missense3D-PTMdb, a dynamic and freely accessible web resource that allows on the fly sequence-structure mapping of missense variants harboured by PTM residues, and by residues in close sequence and structural proximity to PTM residues. Missense3D-PTMdb is freely available for both academic and commercial use and is available at https://missense3d.bc.ic.ac.uk/ptmdb.

## Materials and Methods

### Data collection

20,421 canonical reviewed (Swiss-Prot) human protein sequences, along with their cellular localisation (intracellular, transmembrane, extracellular), were obtained from the UniProt database ^10^. Data on 63 different types of PTMs and the corresponding wild-type residues on which they occur were retrieved from UniProt and from high quality experimentally verified phosphorylation sites identified by Ochoa et al. ^11^ and by Rega et al. ^12^. Phosphorylation sites from Rega et al. were extracted from Supplementary Files S2 and S5, and combined and filtered to include only those with confidently assigned positions within the peptides identified by mass spectrometry. The phosphorylation sites deposited in UniProt are derived from: (i) large-scale proteomics databases, such as PRIDE ^13^ and PTMeXchange ^14^; (ii) literature mining; or (iii) sequence similarity-based prediction. Evidence codes are provided by UniProt to identify the source and confidence level of each PTM annotation.

Data on naturally occurring human missense variants from 1000 Genomes, ExAC/GnomAD, ESP, NCI-TCGA, and COSMIC v71 were obtained from UniProtKB (version June 2025) ^15^ and integrated with variants from the ClinVar database (human built GRCh38) ^16^. Variant clinical significance was also obtained from ClinVar (last updated in June 2025). Variants arising from different genetic changes but resulting in the same amino acid substitution were considered as a single entry, provided they were recognized by the same dbSNP and ClinVar identifiers.

3D structures of human proteins were obtained from the AlphaFold database ^17^. No single AlphaFold 3D model is currently available for proteins longer than 2,700 residues. These proteins were designated as having “no structural data”. For each residue, the AlphaFold pLDDT score, a metric of model confidence, was also retrieved; a pLDDT score ≥ 70 is considered indicative of good structural reliability.

All variants and PTMs were mapped to their corresponding UniProt sequences and available AlphaFold models. In cases where discrepancies in residue numbering occurred between UniProt and AlphaFold sequences, an error message is displayed to notify users. For any queried residue, atomic coordinates are used to calculate pairwise Euclidean distances in 3D space between the target residue and all other residues. Distances are determined by measuring the shortest interatomic distance between residues using coordinate data from the AlphaFold PDB files. These distances are calculated on the fly every time a Results page is loaded on the website.

### Database and web development

All data were stored in a relational database built using MySQL. A dedicated website for querying the database was developed using the Python-based Django framework. Protein structure visualization was implemented using JSmol ^18^.

## Results and Discussion

The Missense3D-PTMdb provides information for 20,421 canonical human proteins, 203,775 PTMs and 11,544,303 human missense variants. Variants are harboured by 7,039,687 variant-harbouring residues (i.e., residues with at least one variant) across 19,264 proteins. There are 199,813 residues that undergo post-translational modifications (PTM residues, i.e., residues with at least one type of PTM) across 15,828 proteins. 20,235 proteins had one single AlphaFold structure model, which allowed mapping of variants and PTMs, where available. It is important to notice that amino acid sequences (motifs), which are crucial for enzyme recognition of residues that undergo PTMs, are not explicitly listed in resources, such as UniProt. Missense3D-PTMdb was developed to cater for the need of users working with variants affecting PTMs and allows easy and friendly sequence-to-structure mapping to visualize variants that occur not only on the PTM residue but also in close sequence and 3D space proximity.

The Missense3D-PTMdb database provides information on 63 distinct PTM types. Phosphorylation sites are the most prevalent, accounting for 130,269 (64%) records, (see Figure 1a), followed by glycosylation, SUMOylation, acetylation, and methylation. The remaining 58 PTM types collectively comprise only 3.4% (n=6951) of all PTM incidences in the database (Figure 1a). It is worth noting that SUMOylation sites were 39,748 (20%) of all PTM records, however this is likely an over-representation due the ongoing migration of high-throughput OTM data in UniProt. We will perform regular updates of our database to align with future UniProt releases.

**Figure 1.**
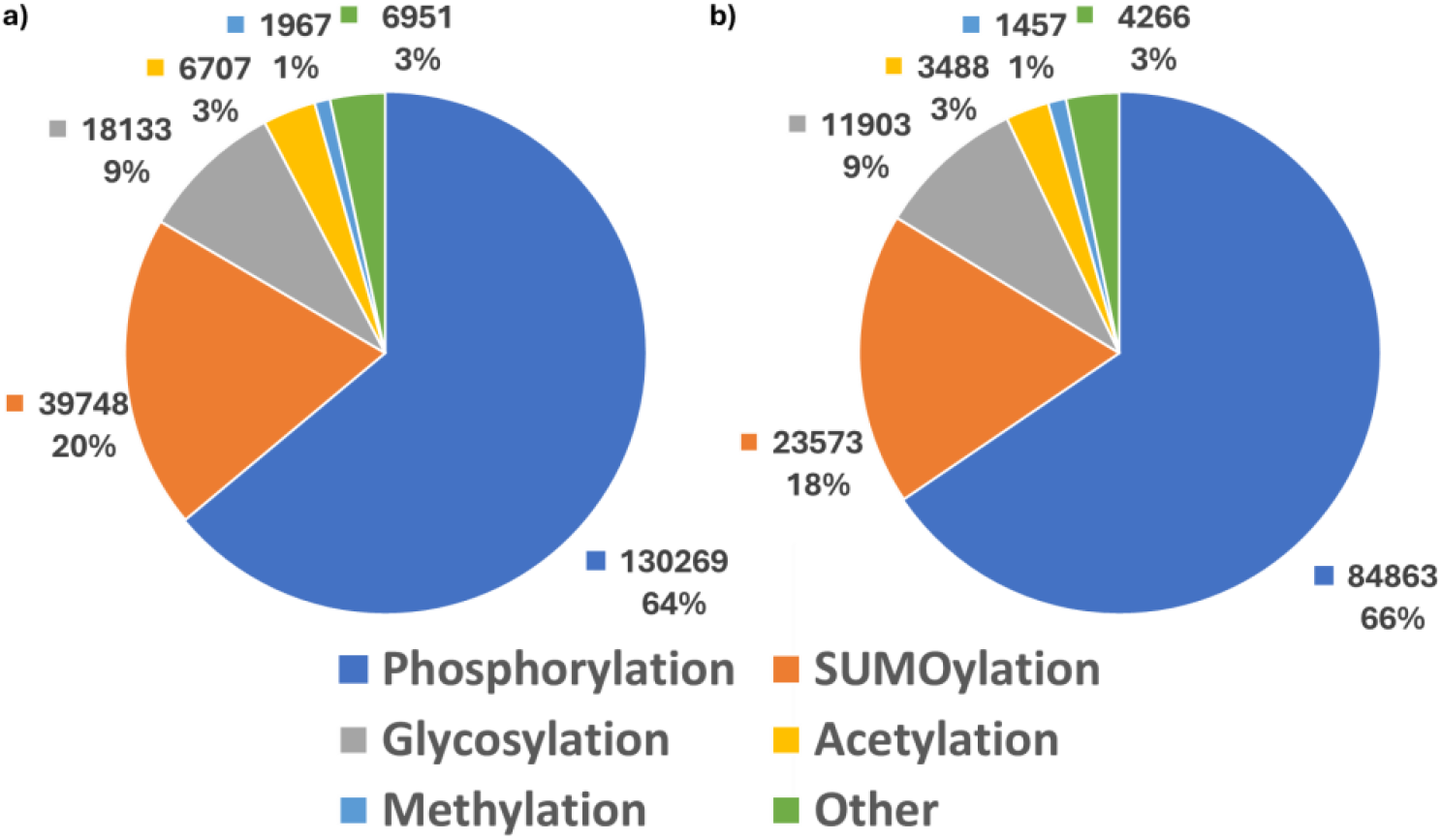
PTMs and variants included in the Missense3D-PTMdb database. a) Overall PTM type distribution; b) PTM residues harbouring at least one missense variant (with or without clinical significance).

### Variants and PTM residues

The Missense3D-PTMdb database includes data on a non-redundant dataset of 11,544,303 naturally occurring missense variants. Of these, 205,645 (1.7%) are harboured by PTM residues. At the time of writing, a total of 571,024 non-redundant variants have clinical annotations from ClinVar and of these 10,510 (1.8%) are on PTM residues.

Overall, 127,106 (64%) PTM residues across 14,014 proteins harbour at least one missense variant. However, this does not consider variants in residues located in close sequence and/or 3D space proximity to a PTM, which can directly or indirectly affect PTMs.

Phosphorylation is the most common PTM type and 84,863 (66%) out of 129,550 variants on PTM residues are on a phosphorylation site (either S, T or Y) (Figure 1b).

### The Missense3D-PTMdb website

The web interface of Missense3D-PTMdb at https://missense3d.bc.ic.ac.uk/ptmdb was designed to be user-friendly for both bioinformaticians and experimental scientists who are non-experts in protein structures, and allows the identification and visualization of PTMs and variants on AlphaFold 3D models. The browser is free to use and requires minimal user input for queries. A valid input consists of: (i) a UniProt ID or gene name for the protein of interest,(ii) the residue position, and amino acid type.

An overview of the Results page is provided in Figure 2. Alongside general information about the query protein (Figure 2a), the Main Results section displays variant and PTM information (Figure 2b), alongside a 3D visualization of the protein’s structure with interactive functionalities (Figure 2c). All information provided in the Results page can be downloaded as an excel file.

**Figure 2.**
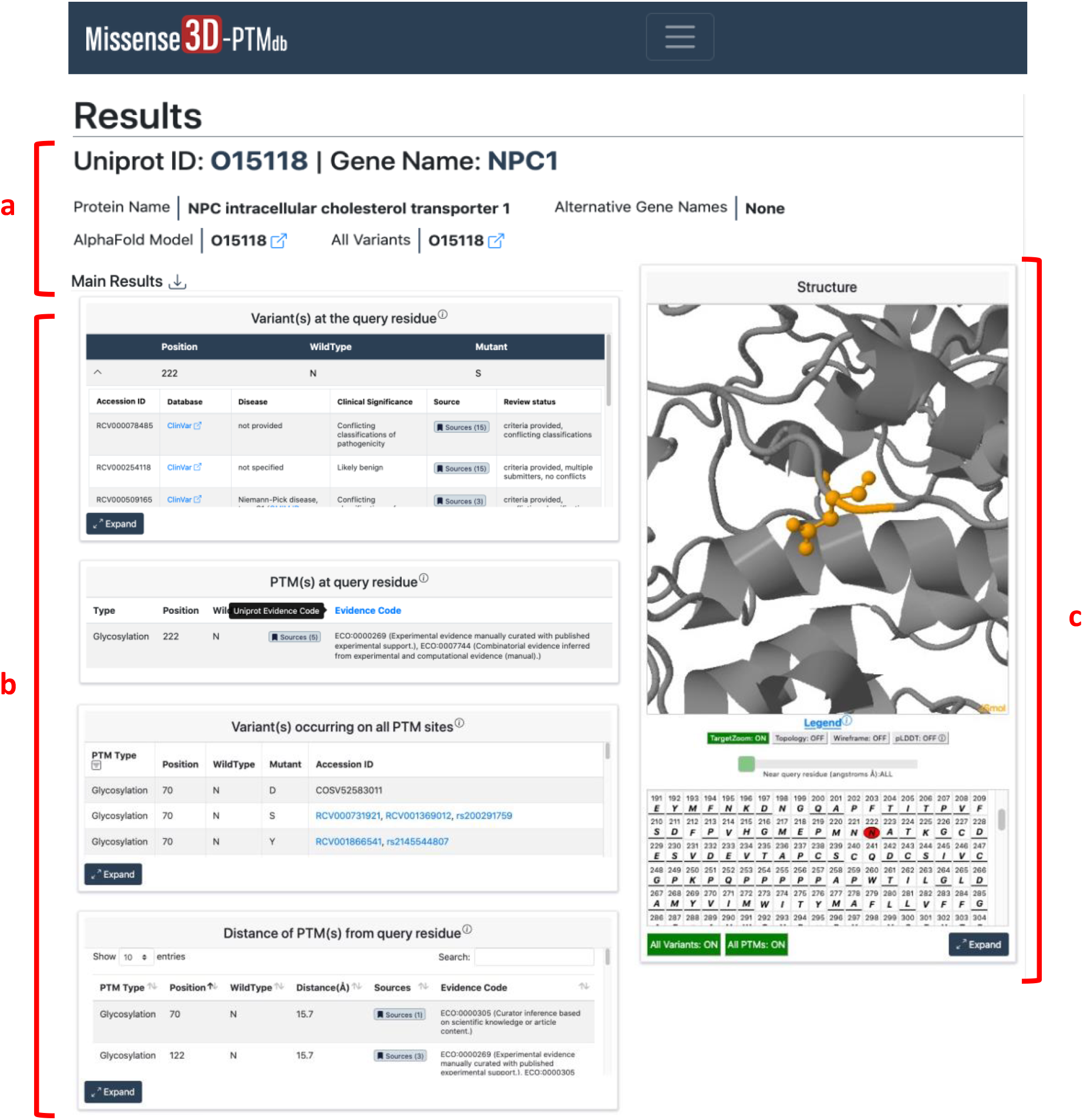
Overview of the Results page. a) Target protein information. b) PTM(s) and variant(s)information at the queried position and on the target protein. c) Structural visualisation of the target protein with the interactive sequence viewer. This example used as query Uniprot ID “P00533”, Position “1104”, and Wild-type “Serine”.

The “Variant(s) at the query residue” table provides a comprehensive summary of known amino acid substitutions harboured by the query residue and their clinical significance when available, whereas the “PTM(s) at query residue” section provides information and source of evidence for PTMs occurring at the query residue (Figure 2b).

To facilitate research on PTM–variant relationship, the “Variant(s) occurring on all PTM sites” section displays information on additional PTMs and their variants identified in the query protein (Figure 2b). Furthermore, to facilitate identification of PTM hot spots, the Euclidean distance (in Angstroms) between the query residue and all other PTMs is provided in the “Distance of PTM(s) from query residue” section. This is a unique feature that demonstrates the strength of our resource and how it can facilitate the exploration of 3D structure data by the non-experts in the field.

### Structural visualization and interactive sequence viewer

The strength of the Missense3D-PTMdb is the sequence-structure mapping and visualization of PTMs and variants using AlphaFold models.

We implemented a sequence viewer that is synchronized with the structural visualization and allows to personalize the 3D structure view. Users can zoom into the query residue and select to display neighbouring residues from 5Å to 30Å in 3D space by using the dedicated “Near query residue (Angstrom, Å)” slider (Figure 3a). Adjusting the distance slider also dynamically updates the sequence viewer, with residues within the selected range highlighted to facilitate identification of nearby residues of interest. Additional residues can be selected on the sequence viewer, and these are automatically highlighted on the corresponding residues in the 3D structure (Figure 3b). Residue marking can be either temporary or persistent, and multiple residues can be highlighted simultaneously. To further enhance usability, the following set of user-controlled options are available (Figure 3c): (i) zoom on/off the target residue, (ii) toggle residue by colouring based on topology (intra, extra and transmembrane) or pLDDT values,(iii)highlight all PTMs and variants.

**Figure 3.**
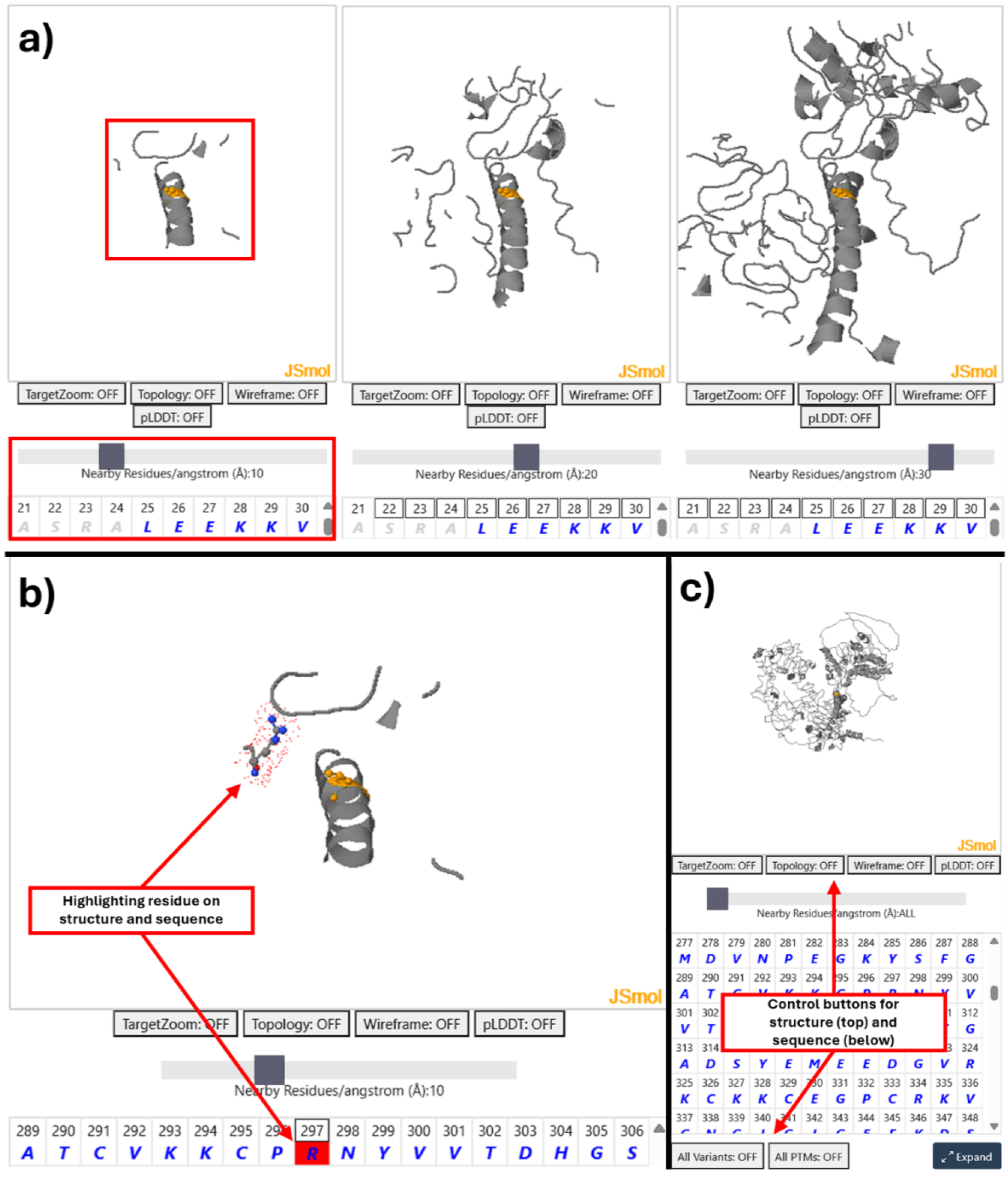
The Missense3D-PTMdb structural visualisation. a) The distance slider at 10 Å, 20 Å, and 30 Å levels from the query residue. b) Highlighting a residue in the 3D structure by selecting it via the sequence viewer. c) Control buttons for both structural visualisation and the sequence viewer.

## Conclusions

In summary, Missense3D-PTMdb is a new freely available resource for the community engaged in understanding the effect of human genetic variation on post-translational modification. The database with its interactive web interface caters for the non-expert user in protein structure data by providing easy mapping and visualization of variants and PTMs onto AlphaFold structures.

## Data Availability Statement

All data included in the Missense3D-PTMdb are openly accessible from resources listed in the Materials and Methods section. All new data generated on the fly are freely available for download from the Results pages.

## Author Contribution Statement

OS, WT, GW, GH, SI developed the database; HZ and RP developed the website; AD designed of the study; HZ and AD wrote the first draft of the paper; all authors approved this manuscript.

## Funding

This study was funded by MRC grant MR/Y031091/1

## Ethical approval

All data used in this project are publicly available. No ethical approval was required for this study.

## Competing interests

The authors declare no competing interests

## Notes

This work was supported by the MRC grant MR/Y031091/1

### Competing Interest Statement

The authors have declared no competing interest.

